# EgGLUT1 is crucial for the viability of larvae of *Echinococcus granulosus sensus lato* by involving its glucose uptake

**DOI:** 10.1101/2021.04.02.438290

**Authors:** Kuerbannisha Amahong, Mingzhi Yan, Jintian Li, Ning Yang, Hui Liu, Xiaojuan Bi, Dominique A. Vuitton, Renyong Lin, Guodong Lü

**Affiliations:** State Key Laboratory of Pathogenesis, Prevention, and Treatment of Central Asian High Incidence Diseases, Clinical Medical Research Institute, The First Affiliated Hospital of Xinjiang Medical University, Urumqi, China; College of Pharmacy, Xinjiang Medical University, Urumqi, China; University Bourgogne Franche-Comté and French National Reference Centre for Echinococcosis, Besançon, France; WHO Collaborating Centre for Prevention and Care Management of Echinococcosis, The First Affiliated Hospital of Xinjiang Medical University, Urumqi, China; Basic Medical College, Xinjiang Medical University, Urumqi, China

**Keywords:** cystic echinococcosis, glucose transporter 1, glucose uptake, WZB117

## Abstract

Cystic echinococcosis (CE) is a zoonotic parasitic disease caused by infection with the larvae of *Echinococcus granulosus sensu lato* (*s.l.*) cluster. It is urgent to identify novel drug targets and develop new drug candidates against CE. Glucose transporter 1 (GLUT1) is mainly responsible for the transmembrane transport of glucose to maintain its constant cellular availability and is a recent research hotspot as a drug target in various diseases. However, presence and role of GLUT1 in *E. granulosus s.l.* (EgGLTU1) was unknown. In this study, we cloned a conserved GLUT1 homology gene (named EgGLUT1-ss) from *E. granulosus sensu stricto* (*s.s.*) and found EgGLUT1-ss was crucial for glucose uptake of the protoscoleces of *E. granulosus s.s..* WZB117, a GLUT1 inhibitor, inhibited glucose uptake of *E. granulosus s.s.* and the viability of the metacestode *in vitro.* In addition, WZB117 showed potent therapeutic activity in *E. granulosus s.s.-*infected mice: a 10 mg/kg dose of WZB117 significantly reduced the number and weight of parasite cysts as well as the reference drug, albendazole. Our data have defined EgGLUT1 as a key *E. granulosus s.l.* vulnerability target, involved in its glucose uptake from the host; this opens a new avenue to identify drugs with an ideal activity profile for the treatment of CE.

## Introduction

Cystic echinococcosis (CE) is a chronic and neglected zoonotic parasitic disease caused by the larvae of the *Echinococcus granulosus sensu lato* (*s.l.*) cluster and listed as one of 17 neglected tropical diseases by the World Health Organization (WHO) (1). CE is distributed worldwide, mainly in South America, Eastern Europe, the Middle East, Russia, and Western China, and the incidence is up to 5% to 10% in highly endemic areas (2, 3). More than 2-3 million cases are estimated worldwide (4). According to the WHO, the costs globally committed for treating CE are more than $3 billion per year (5). China has a high prevalence of human CE, accounting for 40 % of global DALYs lost worldwide (6). Between 2012 and 2016, CE was endemic in 368 counties, with an estimated 166,098 cases of CE in China (7).

The early symptoms of CE are not obvious, and most CE patients are already in advanced stages when they seek medical treatment (2). This treatment mainly includes surgical and anti-parasitic drug treatments, but surgery is not always possible and may be complicated by postoperative recurrence and secondary infection or biliary leakage (8). For some patients who are not suitable for surgical treatment, antiparasitic treatment has become the first choice; in this situation, the germinal layer of the metacestode is the main target of the drug. Anti-parasitic treatment is also used before and after operation to prevent a recurrence, with the possibly spilled protoscoleces (PSCs) as main targets (9). Currently, benzimidazoles, mebendazole and more often albendazole (ABZ), are the only drugs that may be used to treat this infection at its metacestode stage, as recommended by WHO, but such drugs are characterized by poor bioavailability, wide inter-individual variations in blood levels, and occurrence of adverse reactions (10, 11). At present, we are missing anti-CE drugs that can effectively replace ABZ; in addition, the killing potential of ABZ on PSCs is not optimal, and it is delayed (11). Therefore, the identification of new drug targets and the development of new therapeutic molecules are of great significance for the treatment of CE.

The results of transcriptome and whole-genome sequencing show that after entering the intermediate host, the anabolic ability of *E. granulosus s.l.* is severely degraded, while its ability to absorb nutrients is greatly increased, and complete metabolic pathways such as glycolysis, tricarboxylic acid cycle and pentose phosphate cycles function efficiently (12, 13). This suggests that *E. granulosus s.l.* requires nutrients such as glucose from the intermediate host’s environment to meet basic physiological functions such as energy metabolism. If glucose is not transported from the host environment in time or cannot be transported to the parasite, the parasite’s death probably occurs, which is suggested by the presence of specific antibodies to *E. granulosus s.l.* in the serum of patients without lesions in endemic areas, and the observation of abortive forms of disease (14). It is considered that interference with glucose metabolism is one of the mechanisms of action of ABZ (15). However, the precise metabolic chain involved in *E. granulosus s.l.* glucose uptake is unknown.

Glucose transporter 1 (GLUT1) is widely distributed across most cell types and responsible for the transmembrane transport of glucose (16). In recent years, GLUT1 has gradually become a research hotspot in the field of metabolic diseases, cancer, and other diseases, and is considered as an important target for drug development (17–19). Recently, Wei *et al.* found that inhibiting the human GLUT1 in the erythrocytes could alleviate their injury caused by *Plasmodium* spp. Infection (20). For some parasites, such as *Trypanosoma* spp. and *Leishmania* spp., hexose transporters have been reported to be involved in the glucose uptake pathway of the parasite (21, 22). In addition, GLUT1 of *E. multilocularis* was shown to have relatively high glucose transport activity and to be a crucial participant in the parasite glucose metabolism (23). WZB117, which binds to the GLUT1 at the exofacial sugar binding site, is a reversible competitive inhibitor of glucose uptake and exchange glucose transport (24), and has now been shown to have therapeutic effects in cancer (25) and *Plasmodium* infection (20). However, we currently do not know whether GLUT1 can be an important drug target in echinococcosis treatment and whether its inhibitor WZB117 has an anti-cystic echinococcosis effect.

In our study, we tested whether we could affect *E. granulosus s.l.* glucose uptake by targeting EgGLUT1 at the metacestode stage of *E. granulosus s.l.*. We thus cloned a conserved GLUT1 homology gene from *E. granulosus sensu stricto* (*s.s.*), the species most widely responsible for CE cases worldwide, and inhibited EgGLUT1 by WZB117 to study its possible impact on the survival of the metacestode *in vitro* and in an experimental animal model.

## Results

### Cloning, and physiological and biochemical characterization of EgGLUT1-ss

To assess whether GLUT1 gene was present in *E. granulosus s.s*, we successfully cloned a conserved GLUT1 homologous gene from *E. granulosus s.s.*, named EgGLUT1-ss (accession number: MW393831) (Supplementary Figure 1), which consists of 500 amino acids (Figure 1a). Bioinformatics analysis showed that EgGLUT1-ss had one major facilitator superfamily (MFS) domain and 12 transmembrane regions, which were a typical feature of the GLUT gene (Figure 1a). This special structure plays an important role in the transmembrane transport of glucose (26). In addition, we also found that EgGLUT1-ss contained two amidation sites, one glycosylation site, one cAMP- and cGMP-dependent protein kinase phosphorylation site, two casein kinase II phosphorylation sites, five myristoylation sites, and five protein kinase C phosphorylation sites (Figure 1a). Whole sequence alignment analysis showed that the highest overall homologies on the amino acid sequence level could be detected between EgGLUT1-ss and the GLUT1 of *E. multilocularis* (96.81% identical); and GLUT1 orthologs were found in various other species (with a 32.77% to 72.33% homology) (Figure 1a). Phylogenetic analysis revealed that the EgGLUT1-ss gene was generally closely related to parasite species GLUT genes, especially helminths, and was more distantly related to mammalian species such as human and mouse GLUT genes (Figure 1b).

**Figure 1.**
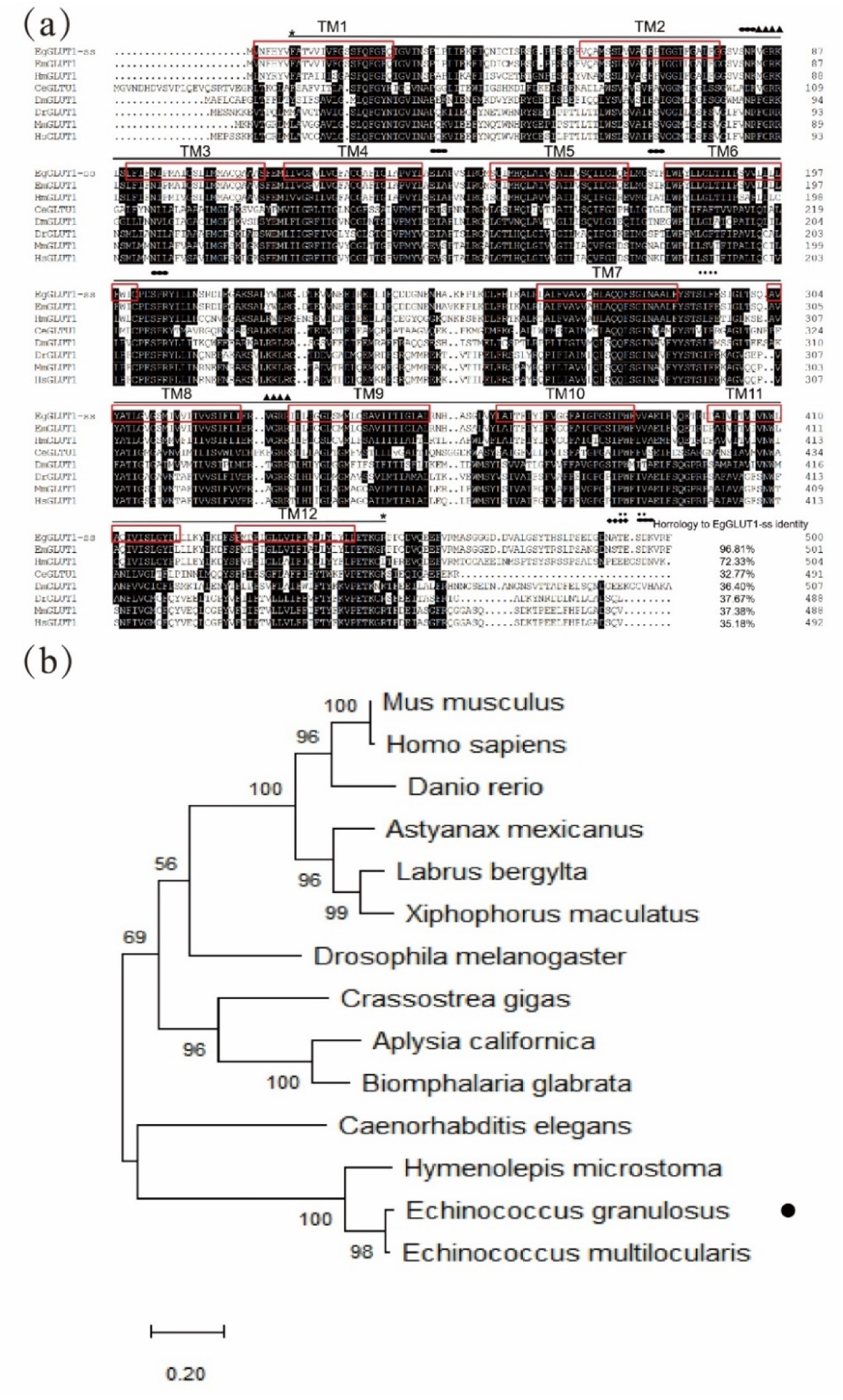
EgGLUT1-ss bioinformatics analysis and its expression level in the metacestode of *E. granulosus s.s.* (a) Multiple-sequence alignments of EgGLUT1-ss from *E. granulosus s.s.* (accession number: MW393831); *Echinococcus multilocularis* (accession number CDS42031.1); *Hymenolepis microstoma* (accession number CDS25463.1); *Caenorhabditis elegans* (accession number NP_493981.1); *Drosophila melanogaster* (accession number NP_001097467.1); *Danio rerio* (accession number NP_001034897.1); *Mus musculus* (accession number XP_006502971.1); and *Homo sapiens* (accession number NP_006507.2). The horizontal line between the two * represents the MFS super family; (▴) denotes the amidation site; (◆) denotes the glycosylation site; (·) denotes Casein kinase II phosphorylation site; (⬬) denotes protein kinase C phosphorylation site; The red box represents the transmembrane (TM) region. (b) Phylogenetic tree constructed using the neighbor-joining method to compare the relationship between EgGLUT1-ss and GLUT1 from other species. The numbers above the branches refer to bootstrap values. The species for sequences included in the phylogenetic analysis are shown behind the branches. EgGLUT1-ss from *E. granulosus s.s.* is indicated by •.

### EgGLUT1-ss Knockdown reduces the viability and glucose uptake of *E. granulosus s.s.* PSCs

To investigate EgGLTU1 function in *E. granulosus s.l.*, we examined the EgGLUT1-ss expression in PSCs and vesicles of *E. granulosus s.s.*. We found EgGLUT1-ss was expressed in both PSCs and vesicles, and EgGLUT1-ss expression level in PSCs was 3.5 times higher than that of vesicles (Figure 2a).

**Figure 2.**
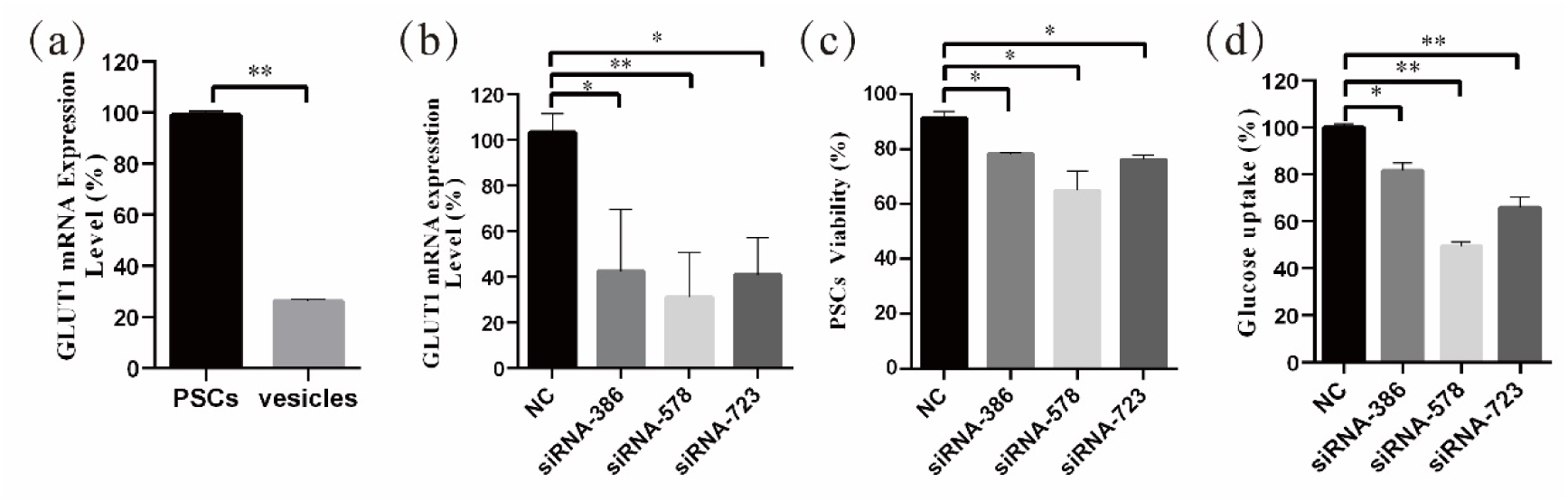
EgGLUT1-ss Knockdown reduces the viability and glucose uptake of PSCs. (a) EgGLUT1-ss mRNA expression in PSCs and vesicles. (b) EgGLUT1-ss mRNA expression at 48 h after siRNA-386/578/723 interference. (c) PSCs viability at 48 h after siRNA-386/578/723 interference. (d) Glucose uptake of PSCs at 48 h after siRNA-386/578/723 interference. NC: negative control group (transfection by independent interference sequence). * and ** indicate that the difference was statistically significant (\**P* < 0.05; \*\**P* < 0.01).

To determine the functional relevance of EgGLUT1-ss in PSCs, we designed three siRNA sequences (siRNA-386/578/723) targeting EgGLUT1-ss. The siRNA-386/578/723 treatment led to a significant reduction in EgGLUT1-ss-mRNA expression of PSCs (Figure 2b) and decreased PSCs viability considerably (Figure 2c). To examine the influence of EgGLUT1-ss knockdown on the glucose uptake of PSCs, siRNA-386/578/723 were transfected into PSCs, and 2 days after transfection, 2-NBDG uptake by PSCs decreased significantly (Figure 2d).

### WZB117 inhibits glucose metabolism and reduces the viability of *E. granulosus s.s.* PSCs *in vitro*

By docking predictions, we found that WZB117 had the potential ability to combine with EgGLUT1-ss. The binding of WZB117 to EgGLUT1-ss involved four hydrogen bonds, with Arg120, Asn29, Asn285 and Gln280. Amino acid residues Phe66, Thr25, Val63, Phe289, Ala286, and Trp410 interacted with the WZB117 molecule through hydrophobic bonds (Figure 3a). After 60 min incubation with WZB117 *in vitro*, the glucose uptake level of PSCs was significantly decreased (Figure 3b). After 48 hours incubation with 25, 50, and 100 μmol/L WZB117 *in vitro*, the glucose content of PSCs was significantly decreased (Figure 3c). In parallel, the ATP content was decreased (Figure 3d) when the PSCs were treated with 50 and 100 μmol/L WZB117.

**Figure 3.**
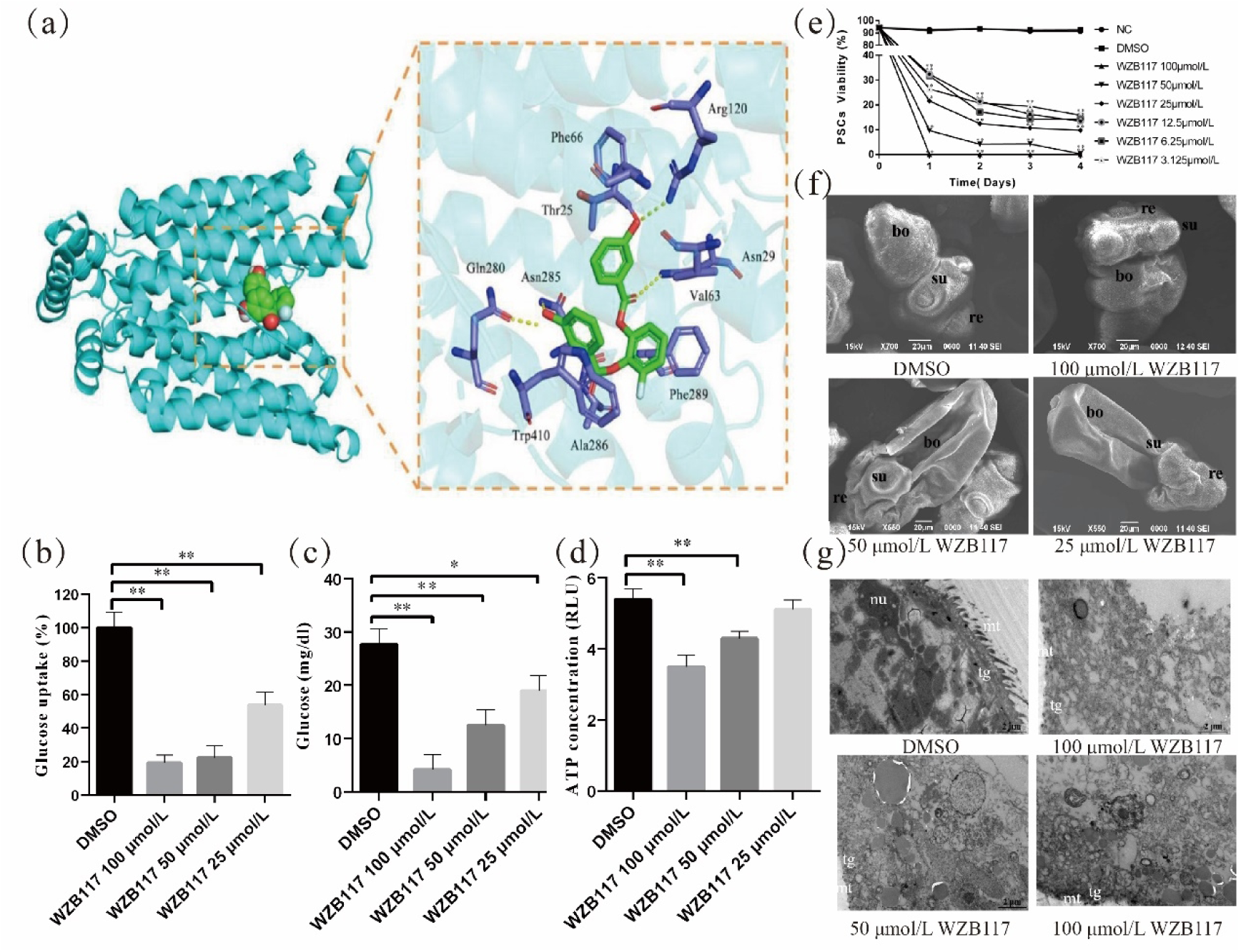
WZB117 inhibits glucose metabolism and reduces the viability of PSCs *in vitro*. (a) Docked structure and interactions of WZB117 binding to EgGLUT1-ss. The image on the left shows that WZB11 binds to the central channel region of EgGLUT1-ss. The image on the right shows the detailed interactions (formation of 4 hydrogen bonds) between WZB117 and amino acid residues of EgGLUT1-ss. (b) Glucose uptake of PSCs after WZB117 incubation for 60 min. (c) Glucose content of PSCs after WZB117 incubation for 48 h. (d) ATP content of PSCs after WZB117 incubation for 48 h. (e) PSCs viability after WZB117 intervention for 4 days. Data are the mean ± S.D. of three independent experiments. (f) Representative SEM images of PSCs after WZB117 incubation for 4 days. PSCs incubated in culture medium containing DMSO served as a control. (g) Representative TEM images of PSCs after WZB117 intervention for 4 days. PSCs incubated in culture medium containing DMSO served as a control. NC: negative control group; DMSO: control group. * and ** indicate that the difference was statistically significant (\**P* < 0.05, \*\**P* < 0.01) compared with control group. re, rostellum; su, suckers; bo, body; mt, microtriches; nu, nucleus; tg, tegument.

To investigate the WZB117 inhibition effect on the viability of *E. granulosus s.s.* larval stages *in vitro*, we analyzed the survival rate of PSCs exposed to various WZB117 concentrations. As shown in Figure 3e, compared to the control group, 4 days-WZB117 exposure led to a dose-dependent decrease in the viability of PSCs in cultures. After exposure to 100 μmol/L WZB117 *in vitro* for one day, all the PSCs died. Scanning electronic microscopy (SEM) showed that, compared to the control group, WZB117-treated PSCs exhibited incomplete rostellum structure, absence of hooks, collapsed suckers and various damages to the other structures of the PSCs, as shown in Figure 3f. Transmission electronic microscopy (TEM) showed that there were significant changes in the ultrastructure of WZB117-treated PSCs: the microtriches disappeared, the nuclei were ruptured, associated with disappearance of the nucleoli, and vacuoles appeared in the cytoplasm. (Figure 3g).

### WZB117 reduces the viability of *E. granulosus s.s.* vesicles *in vitro*

Cysts are the lesions produced by *E. granulosus s.l.* in the intermediate host and their development is related to the pathogenicity of this helminth (2). Therefore, we further explored the effect of WZB117 on such *in vitro*-cultured vesicles exposed to increasing concentrations of WZB117. The viability of the vesicles was significantly decreased when they were exposed to various concentrations of WZB117 for 4 days; it was decreased by 100% on day 3 when exposed to 100 μmol/L WZB117 (Figure 4a). These data and morphological changes indicate that WZB117 had a significant effect on the reduction of the viability of *in vitro-*developed *E. granulosus s.s.* vesicles. Compared with the control group, the SEM features of the vesicles in the WZB117 group changed significantly: the germinal layer was condensed, and the vesicles collapsed. The degree of collapse was positively correlated with the dose of WZB1117 (Figure 4b). TEM images obtained from the WZB117-treated vesicles showed that the structures of germinal layer and laminated layer were damaged, lipid droplet appeared, the microtriches were reduced, nucleoli disappeared, the heterochromatin edges were clustered, and the cortical matrix was fuzzy (Figure 4c).

**Figure 4.**
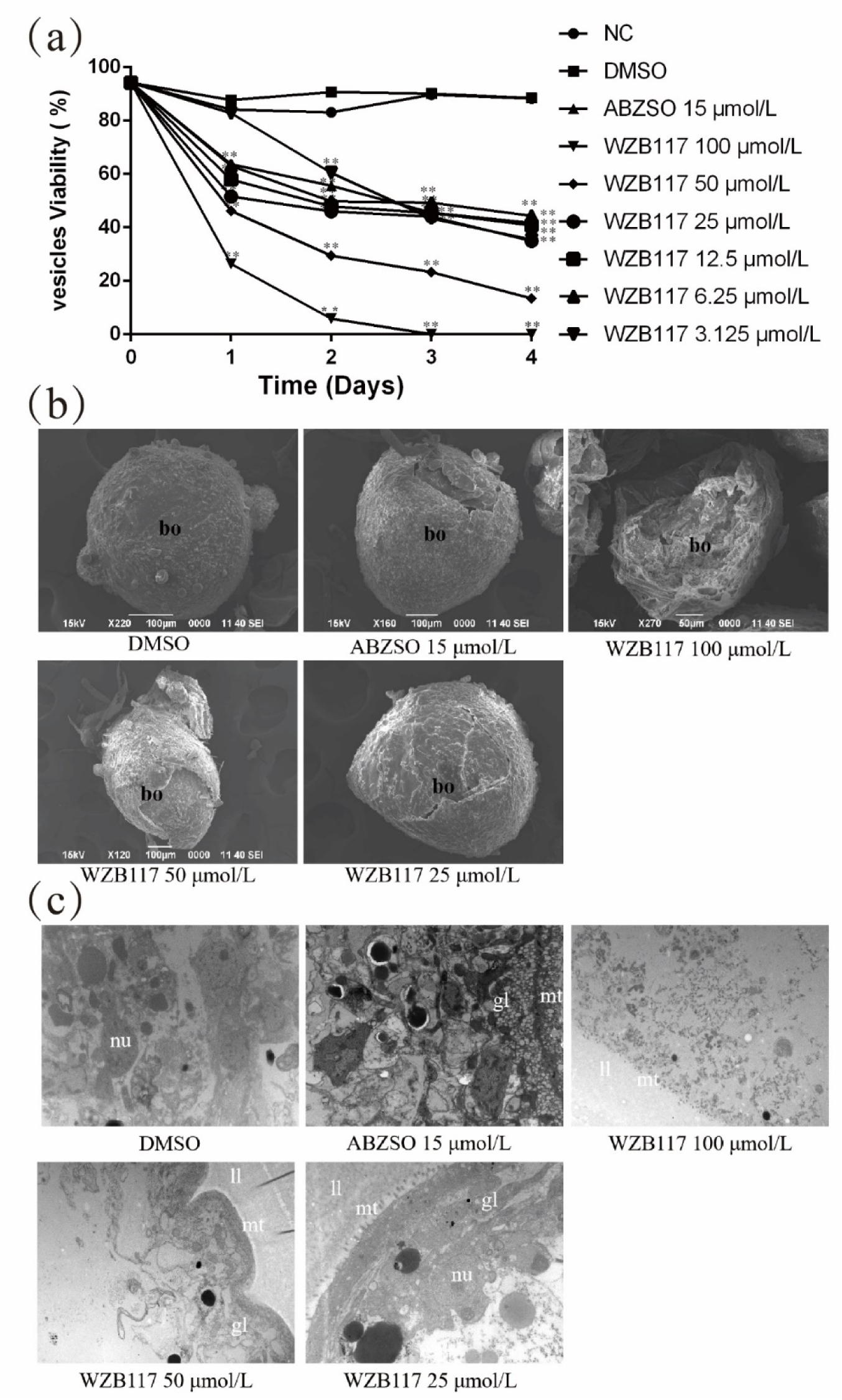
WZB117 reduces the viability of *E. granulosus s.s.* vesicles *in vitro*. (a) Vesicle viability of *E. granulosus s.s.* after WZB117 incubation for 4 days. NC: negative control group; DMSO: control group. * and ** indicate that the difference was statistically significant (\**P* < 0.05, \*\**P* < 0.01) compared with control group. (b) Representative SEM images of vesicles after WZB117 incubation for 4 days. Vesicles incubated in culture medium containing DMSO served as a control. (c) Representative TEM images of vesicles after WZB117 incubation for 4 days. Vesicles incubated in culture medium containing DMSO served as a control. bo, body; gl, germinal layer; ll, laminated layer; mt, microtriches; nu, nucleus.

### WZB117 treatment affects glucose/ATP levels and effectively prevents the growth and of cysts in *E. granulosus s.s.-*infected mice

We measured the glucose and ATP content of the cysts *in vivo*. There was a marked reduction in glucose and ATP levels in the cyst tissue; the reduction was less marked in the cyst fluid (Figure 5a and b). A significant reduction of glucose and ATP levels in the cyst fluid was observed for the 20 mg/kg dose of WZB117 (Figure 5c and d).

**Figure 5.**
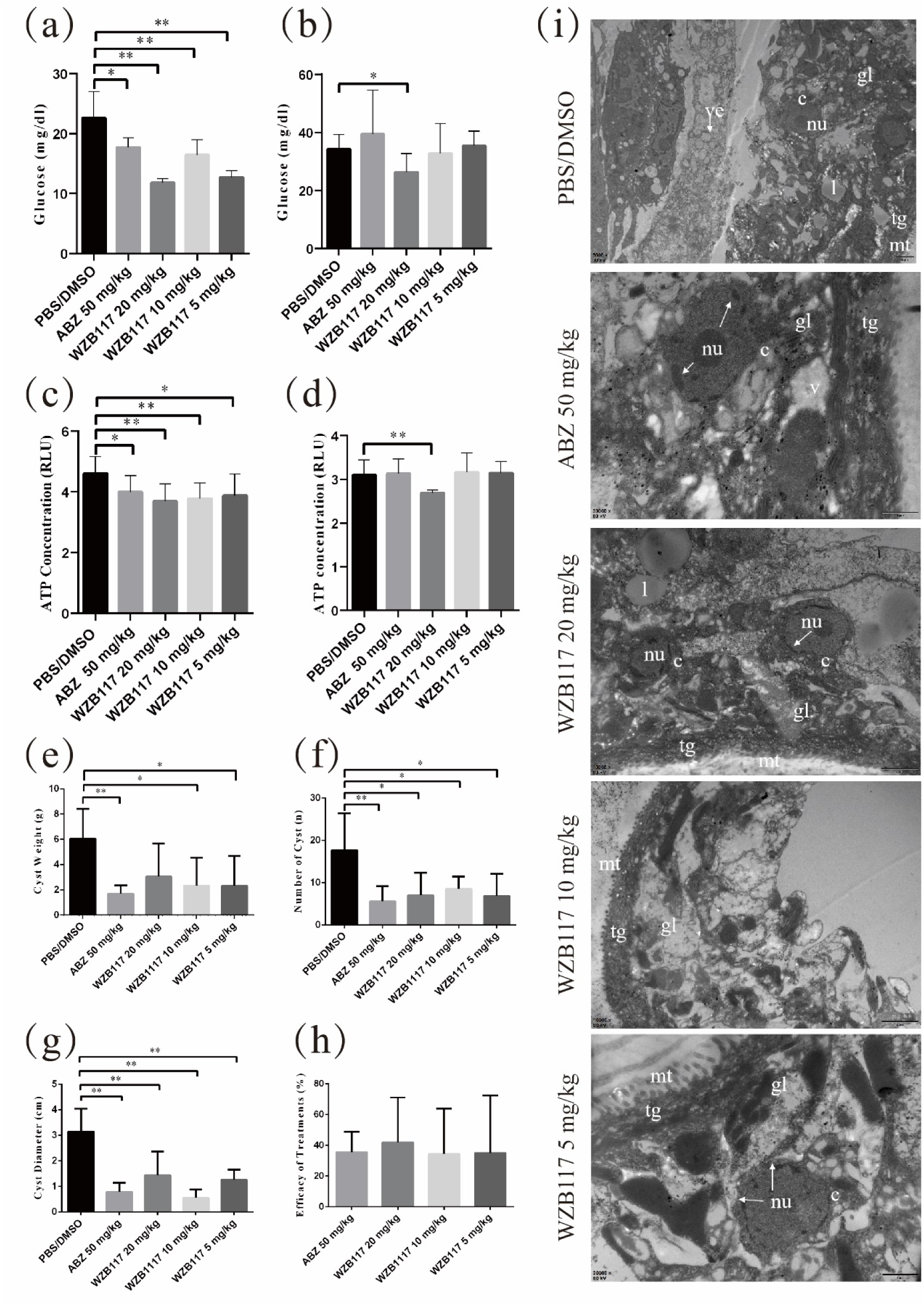
WZB117 treatment effectively inhibits the growth of cysts and reduces glucose and ATP levels in *E. granulosus s.s.-*infected mice. (a) Glucose content of cyst tissue in *E. granulosus s.s.*-infected mice after WZB117 treatment for 28 days. (b) Glucose content in the cyst fluid after WZB117 treatment for 28 days. (c) ATP content in the cyst tissue. (d) ATP content in the cyst fluid. (e) Cyst weight in *E. granulosus s.s.*-infected mice. (f) Cyst number in *E. granulosus s.s.*-infected mice. (g) Cyst diameter in *E. granulosus s.s.*-infected mice. (h) Efficacy of 28-days treatments in *E. granulosus s.s.*-infected mice. (i) Representative TEM images of cysts in *E. granulosus s.s.*-infected mice. PBS/DMSO: control group. * and ** indicate that the difference was statistically significant (\**P* < 0.05, \*\**P* < 0.01) compared with control group. gl, germinal layer; ll, laminated layer; tg, tegument; l, lipid droplet; mt, microtriches; nu, nucleus; c, cell; arrow, broken nucleus.

After 6 months of WZB117 treatment, the weight of cysts recovered from WZB117-treated mice was significantly lower than that recovered from the control group (Figure 5e). Additionally, the number and diameter of cysts in WZB117-treated mice were lower than those of the control group (Figure 5f and g). Overall, the ‘therapeutic’ effect of 10 mg/kg WZB117 was comparable to that of ABZ (Figure 5h).

In the control group (Figure 5i), the cysts had no significant structural abnormalities when observed with TEM. However, the structure of cysts removed from the treated groups was damaged at different degrees (Figure 5i). Compared with the control group, cysts from WZB117-treated mice had loose germinal layers, the microtriches were shortened and reduced dramatically in length, the tegument was thinned, the structure of the laminated layer was destroyed, and the nuclei were condensed.

For a preliminary assessment of the safety of WZB117 in the treatment of CE in experimental hosts, we measured the usual indices of adverse effects. There were no differences in body weight between treated and control mice. The aspartate transaminase (AST) level was significantly lower in the 5 mg/kg WZB117-treated groups than in the control group (Supplementary Figure 2a) and there were no significant differences in alanine aminotransferase (ALT) and alkaline phosphatase (ALP) levels, the most accurate markers of liver damage (Supplementary Figure 2b and c). The 5 mg/kg WZB117-treated group had lower blood urea levels than the control group (Supplementary Figure 2d) but all observed values remained in the physiological range and creatinine levels were similar in all WZB117-treated groups and in the control group (Supplementary Figure 2e). Blood glucose levels were also not different in WZB117-treated mice and in control mice (Supplementary Figure 3). Overall, WZB117, at the dosages used in our study did not cause significant toxicity in mice.

## Discussion

The energy-associated metabolic pathways are essential for the survival of parasites and their adaptation to their successive hosts (27); glucose metabolism is the main energy source (12). Glucose metabolism pathways, especially glycolysis, had been evidenced as a target for anti-echinococcosis drugs (28). However, the upstream pathway of glucose metabolism in *E. granulosus s.l.*, which is involved in glucose uptake from the host, was still elusive. In this study, we identified EgGLUT1-ss as a crucial protein involved in glucose uptake by *E. granulosus s.s.* metacestode and demonstrated that functional EgGLUT1-ss was essential for energy-sourcing and survival in the intermediate host of the parasite. We also characterized EgGLUT1-ss as an important drug target against this larval stage and its inhibitor WZB117 as a candidate anti-CE drug.

Glucose metabolism pathway not only produces ATP for the physiological functions of eucaryote cells, but also provides an important source of carbon to support the biosynthesis of nucleotides and non-essential amino acids (29). *E. granulosus s.l.* produces ATP for its own growth by both aerobic and anaerobic carbohydrate metabolism pathways (30). Tsai *et al.* (12) and Zheng *et al.* (13) found that *E. granulosus s.l.* has complete glucose metabolism pathways, including glycolysis, the tricarboxylic acid cycle, and the pentose phosphate pathway. Wu *et al.* (31) showed that glycolytic enzyme, triosephosphate isomerase, as a drug target for the control of schistosomes (32) and of *Plasmodium* spp. (33), was also involved in the growth and development of *E. granulosus s.l.*. In addition, other glycolytic enzymes, such as 6-phosphofructokinase 1 and pyruvate kinase, have been identified in *E. granulosus s.l.* (34), and Hemer *et al.* (35) showed that the insulin receptor pathway could regulate *E. multilocularis* glucose metabolism. In this study, we found that suppressing the function of EgGLUT1-ss, a member of the glucose transporter 1 family that we cloned from *E. granulosus s.s.,* the species most frequently responsible for CE worldwide, not only inhibited glucose uptake and ATP content by *E. granulosus s.s.*, but also affected the viability of both vesicles and PSCs of *E. granulosus s.s. in vitro* and cysts developed *in vivo*. Glucose, a polar and hydrophilic molecule, cannot pass through hydrophobic cell membranes; GLUT1, which is embedded in the cell membrane, carries out the function of glucose transport to provide glucose supply (36–38). GLUT1-like transporters have been identified in trypanosomes such as *T. brucei, and T. cruzi,* and in *Leishmania* spp. (39), as well as in *E. multilocularis*, another species of the *Echinococcus* genus; in that species, EmGLUT1 has a high glucose transport activity and likely plays an important role in glucose uptake from its host at the larval stage (23). Like *E. multilocularis*, *E. granulosus s.l.* is also dependent on its intermediate host-derived glucose; we may thus suggest that EgGLUT1, a crucial upstream member of glucose metabolism pathways acting as a switch for glucose to enter the parasite, serves as an energy-provider from the host’s glucose to the *E. granulosus s.l.* metacestode, and is thus essential for its growth and survival.

The systemic anti-parasitic treatment of CE currently relies on the continuous administration of either of two benzimidazole carbamates, ABZ and mebendazole, which are the only drugs clinically efficient to interrupt the larval growth of *E. granulosus s.l.* Both drugs interfere with glucose metabolism: their mechanism of action has been associated with a marked inhibition of pyruvate kinase, phosphoenolpyruvate carboxykinase and ATPase (40). In addition, a few bioenergetic modulators have shown significant inhibition of parasite viability in preclinical *in vivo* and *in vitro* experiments. As examples, 3-bromopyruvate which blocks glucose entry into the glycolysis pathway by inhibiting hexokinase (HK) (41) has been shown to inhibit *Echinococcus* spp. viability *in vitro* and *in vivo* (28); tacrolimus, a rapamycin-target protein inhibitor, exerts anti-CE effects *in vivo* and *in vitro* and affects the glucose metabolism of cysts *in vivo* (42); and metformin can reduce the larval viability of *E. granulosus s.l.* by inhibiting its glucose metabolism pathway (43). However, none of these compounds have succeeded in reaching the pre-clinical stage of drug development for CE.

Synthetic GLUT1 inhibitors, such as BAY-876, WZB117 and STF-31, are still in the pre-clinical research stage, but there are obvious experimental data showing that the inhibition of glucose uptake they achieve has a real therapeutic potential (44–46). Previous studies have reported that WZB117 significantly inhibits the proliferation of cancer cells by reducing the transporter function of GLUT1 (47–49) which was overexpressed in cancer cells (50), maybe through the reduction of ATP and glycolytic enzyme levels. Resveratrol, which has a GLUT1 inhibitory effect, has been endorsed by the FDA for the treatment of spinocerebellar ataxia (51). WZB117 was also shown to inhibit the growth of blood-stage *P. berghei* and reduce glucose uptake in the red blood cells by breaking redox balance (20). In our study, we found that GLUT1 inhibitor WZB117 significantly reduced the viability of *E. granulosus s.s..* Our *in vitro* results showed that WZB117 inhibited the function of EgGLUT1-ss, leading to a reduction in glucose and ATP levels, which ultimately led to the death of both the PSCs and metacestode vesicles *in vitro*. In our *in vivo* experiments, we found that WZB117 achieved the same therapeutic results as ABZ, at a lower dose. The lower levels of glucose content in the cyst wall and the cyst fluid we found in the mice treated by WZB117 may suggest that WZB117 could be slightly more effective than ABZ; which may be due to the different targets of the two drugs, inhibition of cytoplasmic and mitochondrial malate dehydrogenase for ABZ (52) on one hand, and EgGLUT1-ss function in glucose uptake and transport for WZB117 on the other hand. The lower anti-CE effect of ABZ may also be related to its poor solubility, thus low cell availability, a well-known disadvantage for its clinical use in the treatment of echinococcosis (8–11). The dual and rapid effect of WZB117 on PSCs and the germinal layer of the metacestode is a supplementary advantage of the GLUT1 inhibitor for its use in CE in the peri-operative period to prevent the development of secondary cysts. Considering the rather frequent adverse effects of ABZ taken at the high dosage necessary to treat CE, and especially its liver toxicity often responsible for drug withdrawal in patients who depend on the drug for their survival (11), WZB117 may likely become an alternative drug to ABZ not only for CE but also AE for which ABZ use is inescapable.

Most of the candidate drugs that were promising, from *in vitro* and experimental studies, against CE and AE have not reached the pre-clinical stage because of adverse effects even superior to those of ABZ (53). Even though it is a promising strategy for the treatment of CE, we cannot ignore the potential adverse effects of WZB117 on the host. It has been previously reported that WZB117 inhibits facilitated glucose transport by competing with sugars for occupancy of the exofacial substrate binding site of the transporter (24). Prolonged inhibition of glucose transport could compromise the normal cellular glycolysis of the host, affect host insulin secretion and cause host’s cerebral energy failure (54). Completing a thorough analysis of all possible adverse events was beyond the scope of this study; however, we conducted a preliminary safety assessment of WZB117 in its use to treat CE, in the murine experimental model. We found no abnormalities in body weight and blood glucose in our mice treated with WZB117 for 28 days. There were no significant changes in the biological parameters under study, especially regarding liver toxicity. Our treatment cycle was only 4 weeks; long-term safety still needs to be further evaluated, since hyperglycemia and lipodystrophy were reported after long-term administration of WZB117 (25); however, our data are very promising to launch pre-clinical studies on peri-operative anti-parasitic treatment of CE, since 1 month is the usual recommended duration of treatment in this situation. The results of our bioinformatics analysis show that the GLUT1 gene sequences of *E. granulosus s.s.* differ significantly from other species, which may true for all species within the *E. granulosus s.l.* cluster but this should be specifically tested whenever the sequences of GLUT1 from other species become available. The design of GLUT1 inhibitors more selective for the protein structure of EgGLUT1 should also be the focus of future developments on CE drugs to avoid interference with the host’s GLUT1 while conserving therapeutic efficacy.

## Materials and methods

### Chemicals

WZB117 and albendazole sulfoxide (ABZSO) were obtained from MedChem Express (USA), and dimethyl sulfoxide (DMSO) and ABZ from Sigma (USA).

### Ethics statement

All procedures carried out with animals were approved by the Ethical Committee of the First Affiliated Hospital of Xinjiang Medical University (IACUC-20130425012). At the end of treatment, all mice were euthanized to avoid the pain of animals to the greatest extent.

### Parasite collection and culture

Sterile PSCs were obtained aseptically from the cysts of sheep infected with *E. granulosus sensu stricto* (*s.s.*) in the Hualing slaughter market of Urumqi in Xinjiang, PR China. All experiments were conducted on this species. Viable and morphologically intact PSCs were cultured using RPMI-1640 medium (Gibco, USA) (10% fetal bovine serum, 100 U/ml penicillin, and 100 μg/ml streptomycin), and maintained at 37°C under a humidified atmosphere containing 5% CO_2_ (55, 56). For *in vitro* experiments on *E. granulosus s.s.* metacestode, *in vitro* sterile cultures were maintained for 4 months to obtain vesicles with a diameter of approximately 2 mm.

### Experimental infection of mice with *E. granulosus s.s.* PSCs

Healthy female Kunming mice (20 ± 2 g of 8 weeks old) were adaptively reared by the Experimental Animal Center of Xinjiang Medical University for one week under controlled laboratory conditions (temperature 20 ± 2°C and 50 ± 5% humidity) (43). The mice were inoculated with 2,000 PSCs in 0.2 mL normal saline by intraperitoneal injection. After 6 months, the mice were examined by B-scan ultrasonography; when the diameter of the lesion was more than 0.5 cm, this indicated that the infection was successful.

### WZB117 treatment *in vitro*

For a first set of experiments, after PSCs were cultured *in vitro* for one week, their viability was tested by 1% eosin staining; to be used in the study, the required percentage of viability required was higher than 90% (57). *In vitro* PSCs treatments were performed with 3.125, 6.25, 12.5, 25, 50 and 100 μmol/L WZB117 (dissolved in DMSO). For a second set of experiments, the PSC-derived vesicles were cultured *in vitro* for four months and then cultured in a 6-well plate (25 vesicles /well). *In vitro* vesicle treatments were performed with 3.125, 6.25, 12.5, 25, 50 and 100 μmol/L WZB117 and 15 μmol/L ABZSO. For both control PSCs and vesicles, the culture medium with an identical amount of DMSO (without inhibitor; final concentration, 1%) was used, and the PSCs and vesicles were cultured in an incubator (5% CO2, 37°C) for 4 days, and each viability test was repeated three times. The collapse of vesicles, loss of swelling and contraction of germinal layer were used as criteria for evaluating vesicle viability (58). Viability of PSCs and vesicles was observed under a microscope every 24 hours (59). Each experiment was performed in triplicate and repeated three times.

### Glucose Uptake Assay

Glucose uptake levels were measured using the 2-[*N*-(7-nitrobenz-2-oxa-1,3-diazol-4-yl) amino]-2-deoxy-D-glucose (2-NBDG) (final concentration, 100 μmol/L) assay. Briefly, the PSCs with siRNA sequence or the WZB117-treated PSCs were incubated at 37℃ for 48 hours. Then, the PSCs were incubated in the darkness with 2-NBDG for 180 min and 60 min at 37°C in 5% CO_2_ humidified atmosphere. The fluorescence intensity was detected at the excitation wavelength of 466 nm and emission wavelength of 540 nm by the fluorescence marker.

### WZB117 treatment *in vivo*

For *in vivo* experiments, WZB117 was dissolved in PBS/DMSO (1:1, v/v) (25). All *E. granulosus s.s.-*infected mice (n = 45) judged appropriate at the 6^th^ month after infection (see above) were randomly divided into five experimental groups: control group (receiving injections of the PBS/DMSO solvent), ABZ (50 mg/kg/day) group (60) and WZB117 (5, 10, 20 mg/kg/day) groups (20). After 28 days of continuous intraperitoneal injection (dosage: 0.1 mL/10 g per mouse), animals were sacrificed by cervical dislocation (57). At necropsy, the peritoneal cavity was opened, the number of cysts was counted, cyst weights and diameters were measured. The efficacy of treatments was calculated as follows: 100 × {(average cyst weight in the control group) - (average cyst weight in the treatment group)} / (average cyst weight in the control group) (57).

### Ultrastructure observations of WZB117 treated PSCs, vesicles, and cysts

After *in vitro* and *in vivo* drug treatment, the PSCs, vesicles, or cysts were fixed with 4% glutaraldehyde for 24 h (28). The samples were processed for SEM using a JEOL1230 (JEOL company, Japan) microscope and TEM using a LEO1430VP (LEO company, Germany) microscopy, as previously described.

### Glucose and ATP content measurements

To determine the glucose and ATP content of cysts in treated mice, the cysts were washed three times with precooled PBS and added with 100 μL of 20 mM Tris buffer (Thermo, USA). The cyst wall was homogenized in a homogenizer for 2∼4 minutes and boiled in a water bath for 5 minutes before centrifugation (at 15,800 g at 4℃ for 30 min) (61). The cyst fluid was processed by gradient dilution according to the instructions in the kit. Glucose Colorimetric Assay Kit (Cayman, USA) was used to determine glucose contents. The ATP content was detected by the ATP Detection Assay Kit (Cayman, USA). Each experiment was repeated three times.

### EgGLUT1-ss cloning

Total RNA was extracted from *E. granulosus s.s.* PSCs by Mini BEST Universal RNA Extraction Kit (Takara, Japan). Me Script II 1^st^ Strand cDNA Synthesis Kit (Takara, Japan) was reverse transcriptionally synthesized for 1^st^ cDNA. The reaction conditions were: incubation at 42℃ for 60 min, heating at 95℃ for 5 min, termination of the reaction, and storage at -20℃ (62). We used the 1^st^ cDNA synthesized by reverse transcription as a template and used the Premix Ex Taq™ Hot Start Version (Takara, Japan) to amplify the full-length sequence of the EgGLUT1-ss gene, primers for the target genes include: EgGLUT1-ss (forward primer 5’-ATGGTTAACTTTCACTACGT-3’ and reverse primer 5’-CTAAAATCTGACCTTATCG-3’). The PCR amplification conditions were: pre-denaturation at 94℃ for 5 min; denaturation at 98℃ for 30 s; annealing at 45℃ for 30 s; extension at 72℃ for 90 s, for a total of 34 cycles. The reaction was terminated after 10 min extension at 72℃; 1% agarose gel was used to determine whether the gene band location was expected. PCR products of EgGLUT1-ss gene were recovered from Agarose Gel with Agarose Gel DNA Extraction Kit (Takara, Japan), and the amplified fragments were cloned into pMD19-T vector with Mighty ta-cloning Reagent Set for Prime STAR (Takara, Japan), and verified by sequencing. The gene was named as EgGLUT1-ss (accession number: MW393831).

### Bioinformatics analysis and construction of the phylogenetic tree of GLUT1

The amino acid sequences of the homologous genes of GLUT1 in *E. multilocularis* (GenBank ID: CDS42031.1), *Hymenolepis microstoma* (GenBank ID: CDS25463.1), *Caenorhabditis elegans* (GenBank ID: NP_493981.1), *Drosophila melanogaster* (GenBank ID: NP_001097467.1), *Labrus bergylta* (GenBank ID: XP_020502389.1), *Astyanax mexicanus* (GenBank ID: XP_007258287.2), *Xiphophorus maculatus* (GenBank ID: XP_023187106.1), *Aplysia californica* (GenBank ID: XP_012944940.1), *Biomphalaria glabrata* (GenBank ID: XP_013087453.1), *Crassostrea gigas* (GenBank ID: XP_019925400.1), *Danio rerio* (GenBank ID: NP_001034897.1), *Mus musculus* (GenBank ID: XP_006502971.1) and *Homo sapiens* (GenBank ID: NP_006507.2) were obtained from GenBank. The post-translational modification sites were predicted using the MotifScan software (63). The transmembrane region was predicted using TMHMM (64). The conserved domain was predicted using GenBank tools. The multiple sequence alignment was analyzed by DNAMAN (Version 7.0.2.176). The phylogenetic tree was constructed by MEGA (Version 10.0.5).

### EgGLUT1-ss-siRNA interference

According to the cloned EgGLUT1-ss Gene sequence (accession number: MW393831), siRNA interference sequences of three EgGLUT1-ss genes were designed using Gene Pharma siRNA Designer 3.0 software. Experimental groups: EgGLUT1-ss-treated groups (siRNA-386/578/723); negative control group (NC, transfection independent interference sequence); untreated groups. The siRNA-386/578/723 were transfected into PSCs cultured *in vitro* by electroporation. The targeting sequence of each siRNA in EgGLUT1-ss cDNA is summarized in Table 1. The EgGLUT1-ss-siRNA interference assay was carried out as previously described (65). Briefly, Electroporation was performed at 125 V using a pulse. For electroporation, 200 μL electroporation buffer (5 mM magnesium chloride, 200 mM glucose, 20 mM Tris, 2 mM 2-Hydroxy-1-ethanethiol, pH adjusted to 7.4 with HCl) containing approximately 4,000 PSCs was placed in a 4-mm cuvette, and the final concentration of siRNA interference sequence was 5 μM. After electroporation, the shock cup was placed in a 37℃ incubator for 30 min and then transferred to a 6-well plate with 2 mL mixed medium for 48 h. After washing with PBS, PSCs in each group were stained with 1% eosin for 5 min, respectively. The viability was calculated on smears by counting using an inverted fluorescence microscope.

**Table 1.**
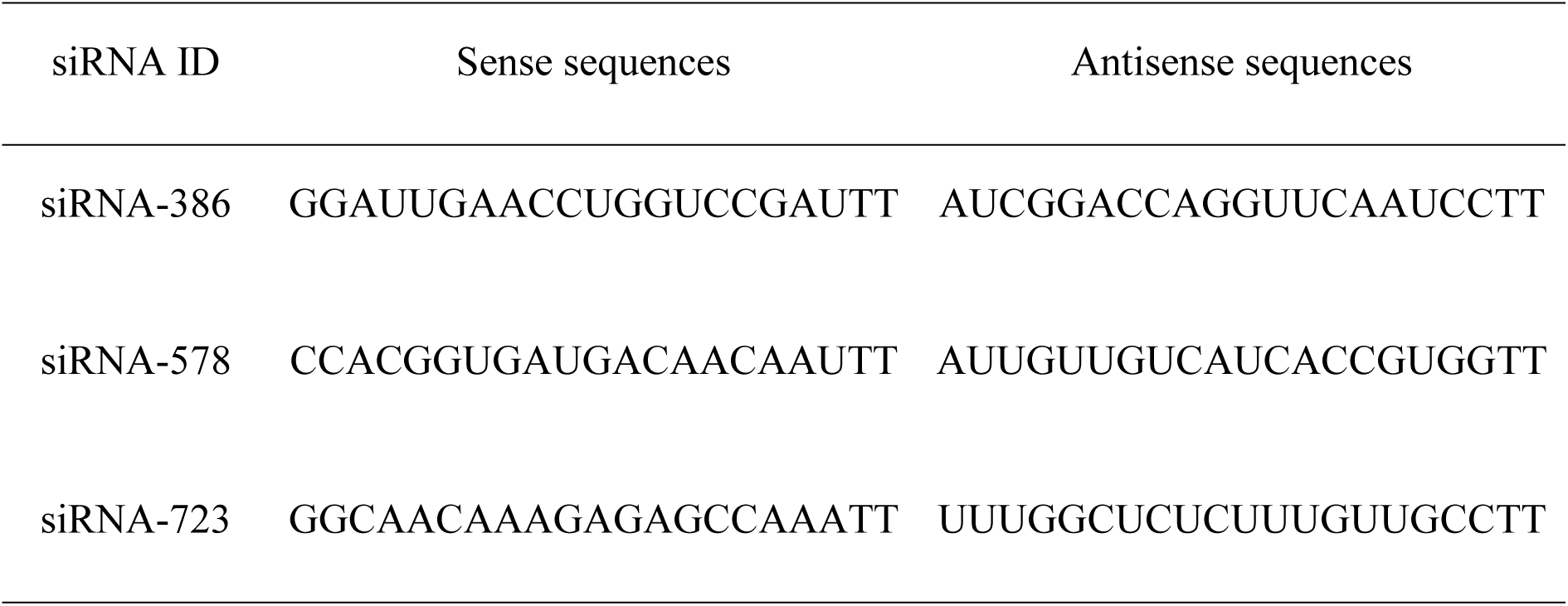
Design of the potential three siRNA sequences targeting EgGLUT1-ss gene

### Quantitative real-time polymerase chain reaction (qRT-PCR) detection

Total RNA was extracted from PSCs and vesicles by Mini BEST Universal RNA Extraction Kit (Takara, Japan), as previously described (66). RNA was converted to cDNA with the PrimeScript™ RT reagent Kit Prime Script TMRT reagent Kit (Takara, Japan), and the cDNA was analyzed by qRT-PCR using TB Green^®^ Premix Ex Taq^TM^ II (Takara, Japan). The primer sequences are shown in Table 2. All samples were run in triplicate using the following cycle parameters: 95℃ for 30 s; 40 cycles at 95℃ for 5 s and 55℃ for 30 s; 72℃ for 1 min; from 95℃, the temperature dropped to 65℃ at a rate of 20.0℃/s. After incubation for 15 s at 65℃, the temperature was increased to 95℃ at a rate of 20.0℃/s. All data were used for standard curve analysis.

**Table 2.**
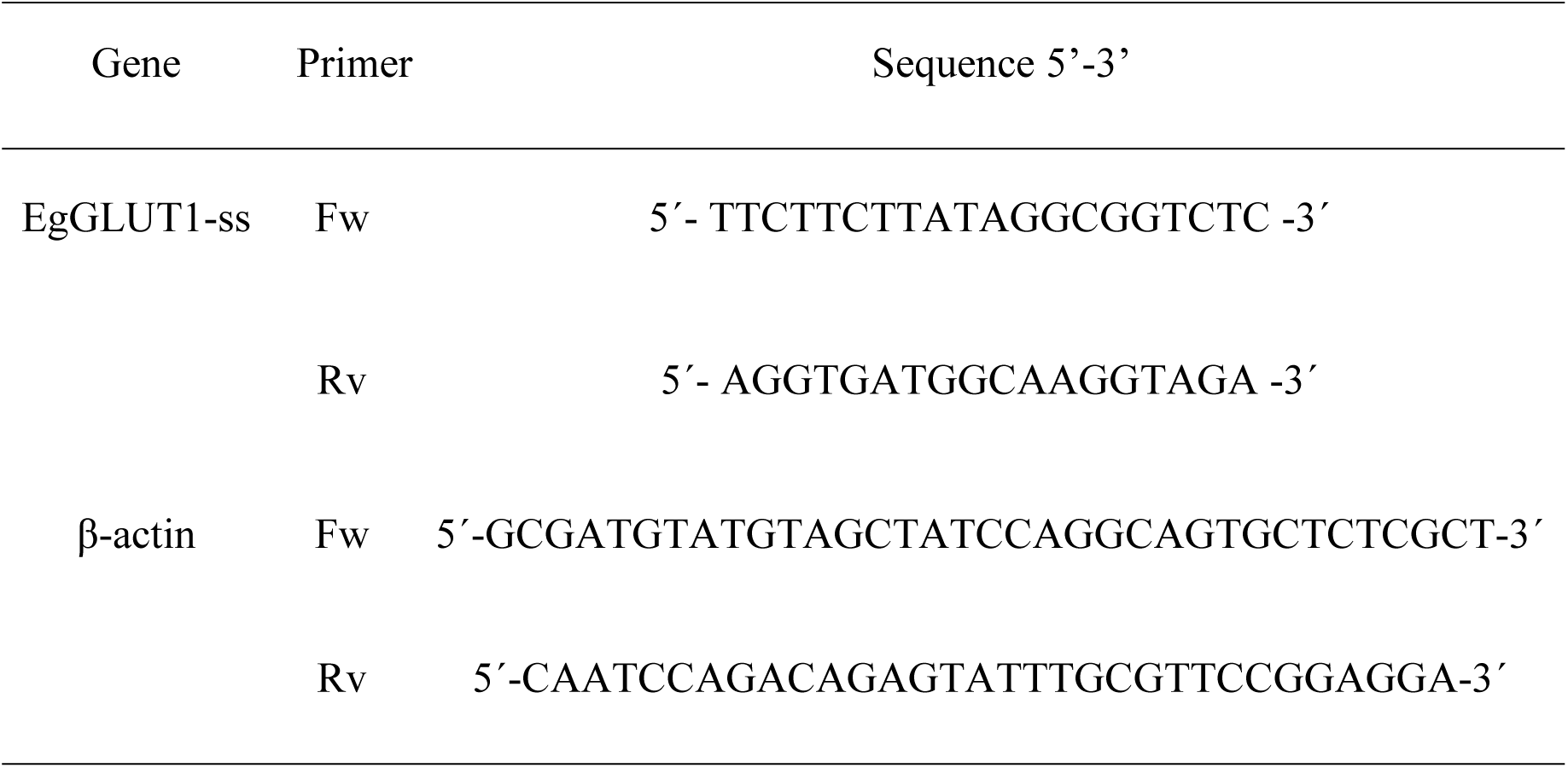
EgGLUT1-ss Gene primer sequences

### Docking study

Through the Pubchem website (https://pubchem.ncbi.nlm.nih.gov/) to retrieve the 3D structure of WZB117, we adopted the Autodock software (version 4.2.6) to hydrogenation processing, calculating charge EgGLUT1-ss protein, and setting the receptor proteins docking lattice parameter. We used AutoDock Vina (version 1.1.2) to dock EgGLUT1-ss and WZB117 and obtained 20 conformations. EgGLUT1-ss and WZB117 were used to chart the binding sites and amino acid residues and PyMOL (version 2.4.0) was used as a 3D diagram to show the interaction between receptor proteins and ligand small molecules.

### Statistical analysis

For the comparison between experimental and control data, Student’s t-test and chi-square test were used to determine its significance. All data were expressed as (arithmetic mean ± standard deviation), and *P* values are indicated in each assay (\**P* < 0.05, \*\**P* < 0.01). All analyses were performed using IBM SPSS Statistics 20 software.

### Linguistic statement

The terminology proposed by the World Association of Echinococcosis was followed all along the text of the publication. Especially, we followed the distinction between ‘vesicles’, obtained *in vitro*, and ‘cysts’ obtained from *in vivo* experiments, and between *E. granulosus sensu lato* for the cluster of species previously known as ‘*E. granulosus’* and ‘*E. granulosus sensu stricto’* for the species used in our experiments (67).

## Acknowledgements

This research was supported by the National Natural Science Foundation of China (NSFC) (82060373, 81760369 and 81360251); State Key Laboratory of Pathogenesis, Prevention and Treatment of Central Asian High Incidence Diseases Fund (SKL-HIDCA-2020-BC3); “Tianshan Cedar” Science and Technology Innovation Talents Support Plan of Xinjiang Uygur Autonomous Region (No. 2019XS13). The funders had no role in study design, data collection and interpretation, or the decision to submit the work for publication.

We thank Zhiqiang Li of the Animal Experiment Center of Xinjiang Medical University for providing help with animal experiments and Haiyan Ren of the Department of Electron Microscopy of Xinjiang Medical University for providing technical support in the use of electron microscopes.

## Transparency declarations

None to declare.

